# Integrative Metagenomic and Blood Biomarker Analysis Reveals Distinct Metabolic Responses to Aerobic and Anaerobic Interventions in Athletic and Non-Athletic Individuals

**DOI:** 10.1101/2024.11.19.624295

**Authors:** Kinga Zielińska, Monika Michałowska-Sawczyn, Tomasz Kosciolek, Paweł P. Łabaj, Barkin Bicakci, Patrizia Proia, Paweł Cięszczyk, Andrzej Kochanowicz, Jan Mieszkowski, Kinga Humińska-Lisowska

**Affiliations:** Malopolska Centre of Biotechnology, Jagiellonian University, Krakow, Poland; Faculty of Physical Culture, Gdansk University of Physical Education and Sport, Gdansk, Poland; Sano Centre for Computational Medicine, Krakow, Poland; Faculty of Health Sciences, University of Lomza, Lomza, Poland; Sport and Exercise Sciences Research Unit, Department of Psychology, Educational Science and Human Movement, University of Palermo, Palermo, Italy

**Keywords:** gut microbiome, exercise-induced physiological adaptation, inflammatory cytokines, microbiome-host interactions, physical activity, shotgun metagenomics, sport

## Abstract

**Background:** The gut microbiome influences physiological responses to exercise through interactions with inflammatory markers and metabolite production. Athletes often exhibit a more diverse gut microbiome that is associated with improved performance, but the mechanisms, particularly in relation to different training modalities, are not fully understood. This study aims to integrate blood serum markers with metagenomic data to explore the interplay between exercise type, gut microbiome, and biochemical responses in athletic and non-athletic individuals.

**Methods:** Fifty-two male participants (endurance athletes, strength athletes, and controls) underwent two maximal exercise tests: the anaerobic Wingate test and the aerobic Bruce treadmill test. Blood and stool samples were collected at multiple time points for biochemical analysis and metagenomic sequencing. Blood markers were correlated with shifts in microbiome composition and function.

**Results:** While most biochemical parameters showed similar trends across all groups post-exercise, SPARC and adiponectin levels showed distinct responses depending on the exercise modality. The strength group showed unique microbiome associations with blood markers after the Wingate test. Baseline enrichment of specific bacteria (*Clostridium phoceensis* and *Catenibacterium spp*.) was associated with an inhibited response to the Bruce test in strength athletes.

**Conclusions:** The integration of metagenomic and blood serum analyses reveals that exercise modality and training background elicit complex physiological and biochemical responses mediated by the gut microbiome. These findings suggest that specific microbial species may play significant roles in recovery and adaptation processes following acute exercise. Further research with larger cohorts is needed to validate these findings and explore microbiome-targeted interventions to improve performance and recovery.

## Introduction

The human gut microbiome is a community of microorganisms, including bacteria, which interact with each other and with their host in a specific ecological niche. The microbiome is a dynamic community, displaying a wide range of functions and producing various metabolites [1] . The human gut microbiome is influenced by various host factors such as sleep [2], diet [3], age [4] and exercise [5]. Due to its ability to adapt to different environmental conditions, it has been extensively studied in relation to sports performance [6].

Over 10 years ago, it was already acknowledged that athletes have a more diverse and richer gut microbiome compared to non-athletes. [7]. This diversity is linked to improved metabolic health, enhanced immunity, and better overall performance. While a number of training-beneficial species and mechanisms through which the gut microbiome interacts with the host in the context of sports are known, many findings remain ambiguous [8].

Exercise, both aerobic and anaerobic, induces specific physiological responses and adaptations reflected in changes in blood serum markers. Aerobic exercise, characterized by sustained, moderate-intensity activities relying on oxidative metabolism, improves cardiovascular efficiency and endurance performance [9]. Anaerobic exercise involves high-intensity, short-duration activities that rely on anaerobic energy pathways and result in increased muscle strength and power [10]. These training modalities influence the secretion of various proteins affecting human organs, including inflammatory markers, hormones, and cytokines [11]. The intensity, direction, and rate of these responses depend on many factors and do not always result in beneficial changes in body structure. Prolonged exercise duration and intensity contribute to increased oxidative stress, which can lead to a higher risk of injury and dysfunction of working tissues [12].

The gut microbiome plays a significant role in modulating these physiological responses through its interactions with inflammatory markers. Specific gut bacteria can influence systemic inflammation by producing metabolites such as short-chain fatty acids (SCFAs), which have anti-inflammatory properties [13]. Conversely, inflammation can alter gut microbiome composition, creating a bidirectional relationship between the microbiome and the host’s immune system [14]. Understanding these interactions is crucial, as they may impact recovery processes and overall athletic performance.

In our previous study, we sought to evaluate the gut microbiome and its response to acute exercise interventions performed at maximal intensity [15]. We found that individuals with an athlete-level VO_2max_ experienced an enrichment of *Bifidobacterium longum*, a probiotic species commonly represented in commercial products, as well as a short-chain fatty acids producer *Roseburia inulinivorans*, after the aerobic effort. In addition, we identified correlations between maximal power output and SCFA producers *Eubacterium rectale, Blautia wexlerae* and *Intestinimonas timonensis*. Various individual responses were observed, however, suggesting complex microbiome interactions.

Here, motivated by numerous studies indicating significant changes in inflammatory markers induced by maximal exercise, we integrate blood serum markers with metagenomic data (originally analysed in our previous study). We begin by outlining markers which change independently of exercise modality and training background. Afterwards, we discuss markers which have similar trends in all participant groups but differ as a result of different exercise modalities. Finally, we determine markers which distinguish the participants based on their training background. We explore microbial associations with the blood markers to identify potential mechanisms driving different responses to training interventions in the control, endurance and strength groups. This allows us to identify baseline microbiome features in participants who experience an excessive stress response to a subsequent intervention.

## Materials and Methods

### 2.1. Experimental Overview

This study is an extension of our previous investigation [15], which aimed to evaluate the gut microbiome and its response to different exercise modalities in different athletic populations. This current study integrates biochemical analyses and advanced statistical and bioinformatic methods to further understand the interplay between exercise, microbiome and biochemical markers.

This is an interventional, case-control, repeated-measures study, focusing on the gut microbiome composition and biochemical responses before and after two different exercise tests: (1) a repeated 30-s all-out Wingate test (WT) and (2) the Bruce Treadmill Test. The fitness tests were performed at the University of Physical Education in Gdansk (Gdansk, Poland) in October 2018, following the same protocol as previously described [15]. The Wingate and Bruce treadmill tests were separated by a 14-day break, during which time participants maintained their regular physical activities and training routines.

### 2.2. Participants

A total of 52 male participants were recruited and divided into three groups: endurance athletes (n=15), strength athletes (n=16), and a control group (n=21). Endurance athletes had at least 5 years of training in race walking, long-distance running, or ski running, while strength athletes had a similar duration of training in weightlifting, powerlifting, or bodybuilding. The control group consisted of physically active men who did not participate in organized training but were involved in recreational sports and attended physical education classes at the University of Physical Education and Sport in Gdansk, Poland. There were no statistically significant morphological differences between the control, endurance, and strength groups. Inclusion and exclusion criteria were consistent with those from our previous study [15].

The study protocols were conducted in accordance with the Declaration of Helsinki and approved by the Bioethics Committee for Clinical Research of the Regional Medical Society in Gdansk (KB-27/18). All participants provided written informed consent. Personal data were encoded to ensure anonymity and privacy.

### 2.3. Measurement of Anaerobic and Aerobic Fitness Level

The following procedures were described in our previous study [15]. In brief, the Wingate Test (WT) was performed on a Monark 894E friction-loaded cycle ergometer (Monark 894E, Peak Bike from Sweden) after a 5min warm-up at 60 rpm (1W/kg). Participants performed two 30-second bouts of maximal effort pedalling against a resistance load of 75 g/kg body mass, with a 30-second rest between bouts, as recommended by Bar-Or [16]. Performance measures included peak power, relative peak power, mean power, and relative mean power. The Bruce Treadmill Test was performed using the Bruce protocol on an H/P/Cosmos electric treadmill [17]. Participants completed a 5-minute warm-up at 60% HR max, followed by running through 10 stages of increasing difficulty, starting at a 10% incline and 2.7 km/h, and ending at a 28% incline and 12.07 km/h. The test continued until the participants became fatigued. Maximal oxygen uptake was measured using a Quark CPET pulmonary gas exchange analyser.

### 2.4. Sample Collection and Measurements of Selected Markers

Procedures were adapted from our previous studies [18],[15]. Blood samples (9 mL) were collected at five time points: immediately before, immediately after, 2 hours, 6 hours, and 24 hours after each test. Venous blood samples were collected in Sarstedt S-Monovette tubes (S-Monovette^®^, Sarstedt AG&Co, Nümbrecht, Germany) equipped with a coagulation accelerator for serum separation. Serum was processed according to standard laboratory protocols, aliquoted at 500 μL, and stored at -80°C until further analysis (up to 6 months). The following markers were measured: adiponectin, follistatin-like 1, IL-1 alpha, IL-6, IL-10, IL-15, leptin, leukemia inhibitory factor (LIF), oncostatin M (OSM), resistin, SPARC, TIMP-1, and transferrin receptor (TfR). Analyses were performed using a MAGPIX fluorescence detection system (Luminex Corp., Austin, TX, USA) with Luminex assays (Luminex Corp.; Human Magnetic Luminex Assay (13-plex)).

In addition, fresh stool samples were collected from all participants at three time points: T0 (W0 or B0), the morning after an overnight fast of approximately 8 hours before the exercise test; T1 (W1 or B1), the same day after the exercise test; and T2 (W2 or B2), the next morning on an empty stomach. Samples were placed in stool containers, immediately placed in thermal bags with cooling inserts, and kept refrigerated for a short time. Samples were delivered to the laboratory within a few hours and immediately snap frozen at -80°C.

### 2.5. Metagenomic sample preparation

The procedures for sample preparation, DNA isolation, quantification, and library preparation followed the methods described in our previous publication [15]. Briefly, DNA was extracted from 200 mg of the frozen stool samples using a modified NucleoSpin^®^ DNA Stool Kit (Macherey-Nagel, Germany) with physical, mechanical, and chemical lysis. DNA concentration and purity were measured using a NanoDrop spectrophotometer and a Qubit fluorometer. Library preparation followed the KAPA HyperPlus protocol and sequencing was performed on an Illumina NovaSeq platform, generating approximately 22 million 150 bp paired-end reads per library. The raw fastq files are openly available in the in the European Nucleotide Archive (ENA) at https://www.ebi.ac.uk/ena/, reference number PRJEB60692.

### 2.6. Statistical and bioinformatic analysis

The raw fastq sequences underwent quality control using TrimGalore [19]. Taxonomic and functional profiles were calculated using MetaPhlAn 4.0 and HumanN 3.7 [20]. The differences between blood serum marker levels at consecutive timepoints were evaluated using the independent T-test. The correlations of the blood markers and metagenomic features were performed in two ways: a) the correlations at single timepoints and b) the changes in blood markers and metagenomic features between two timepoints, correlated using Spearman with Benjamini-Hochberg correction for multiple testing [21]. Finally, we used Maaslin2 with default settings to perform differential enrichment analysis of species and pathways distinguishing the baseline microbiomes of the responders and nonresponders to the Bruce intervention [22].

## Results

### Most biochemical changes are independent of training background and exercise modality

A previously published by us cohort consisting of 52 male participants was divided based on training background into endurance (n=15), strength (n=16) and control (n=21) groups [15]. All participants were subjected to two maximal effort interventions: an anaerobic-focused Wingate test and an aerobic-focused Bruce test. Stool samples were collected before and twice after each intervention. For the purpose of this study, we included biochemical parameter collection timepoints that approximated the time of stool collection (there were three stool and five biochemical collections in total).

Most biochemical parameters showed similar trends in all groups after the Wingate (Figure 1 top) and Bruce (Figure 1 bottom) interventions. Oncostatin M (OSM) and IL-15 levels (Figure 1a, b) showed significant decreases after both the Wingate and Bruce tests in all groups (endurance, strength, and control) at 24 hours post-exercise. Follistatin-like 1 (FSTL1) levels (Figure 1c) showed a slight increase 6 hours after the Wingate test and a statistically significant decrease in the strength group 24 hours after the Wingate test (p < 0.05). After the Bruce test, FSTL1 levels also showed a slight increase at 6 hours followed by a statistically significant decrease at 24 hours in all groups (p < 0.05), suggesting a consistent pattern of initial increase and subsequent decrease in response to both types of exercise. A statistically significant decrease in leptin levels was observed 24 hours after the Wingate test in the strength group (Figure 1d, p < 0.0001). While endurance and control groups did not display statistically significant trends, they would decrease as well. Conversely, levels of leukemia inhibitory factor (LIF), resistin, interleukin-6 (IL-6), and transferrin receptor (TfR) (Figures 1e-h) remained stable in both the Wingate and Bruce tests. These markers did not show significant changes, indicating that they were not significantly affected by the exercise interventions. In conclusion, even if significant changes were observed only after one test, they would follow a similar pattern after the other intervention. Therefore, the aforementioned parameters were not associated with the differential responses to exercise modalities among the groups.

**Figure 1.**
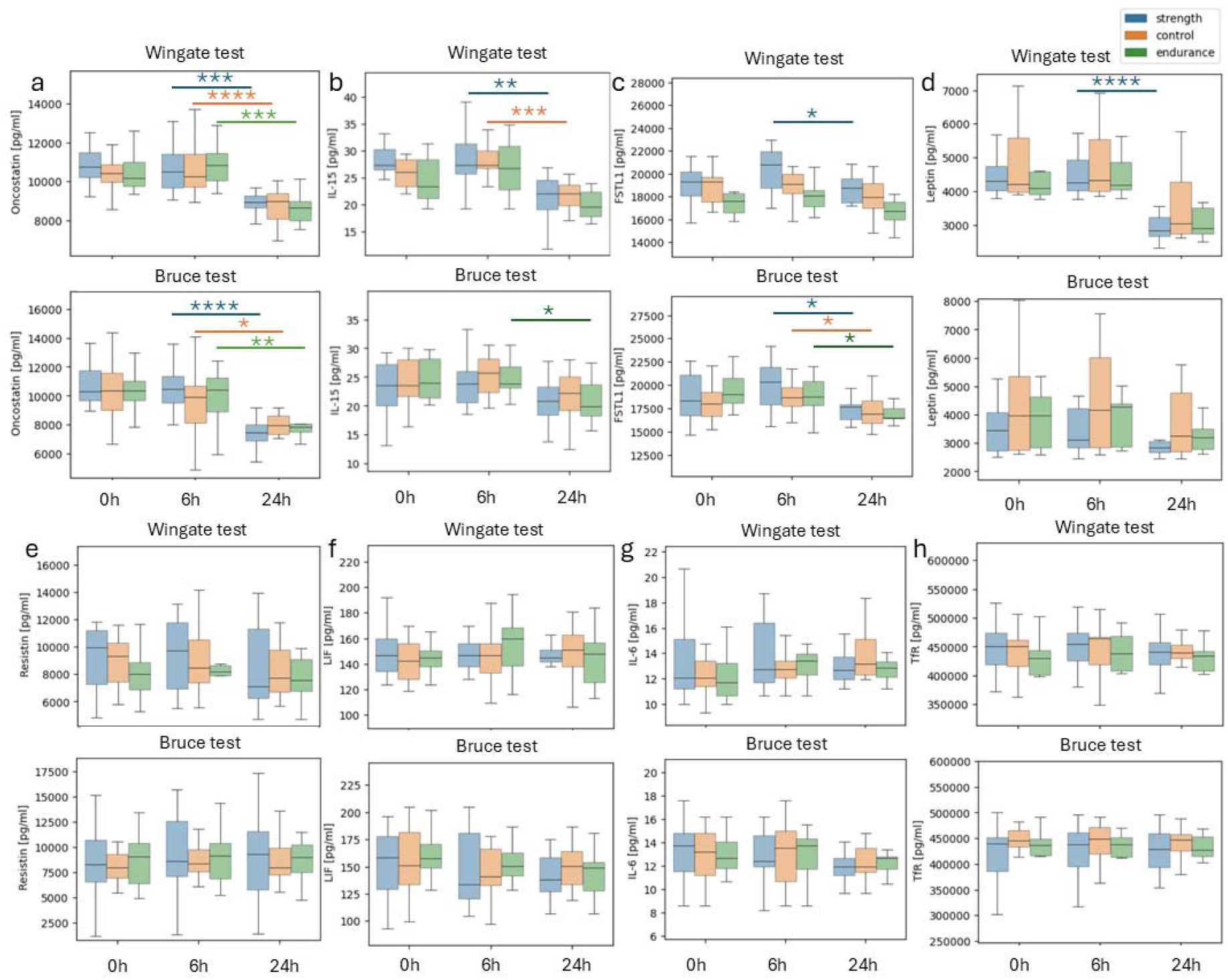
Post-exercise changes in biochemical markers in all groups following the Wingate (top) and Bruce (bottom) tests. Most biochemical parameters show similar trends in all groups after the Wingate (top) and Bruce (bottom) tests. Blue: strength, orange: control, green: endurance. a – oncostatin, b – IL-15 (interleukin 15), c – FSTL1 (Follistatin-like 1), d – leptin, e – resistin, f – LIF – leukemia inhibitory factor, g - IL-6 (interleukin 6), h - TfR (transferrin receptor); ^*^=p<0,05, ^**^=p<0,01, ^***^=p < 0.001, ^****^=p < 0.0001.

### Nucleotide recycling as a modulator of responses to different training efforts

SPARC and adiponectin were the only biochemical parameters which behaved differently as a result of the interventions, similarly in every group (Figure 2 a-d). SPARC levels showed a significant increase only in the control group 24 hours after the Wingate test (p < 0.05). Although there was an unsignificant trend in the endurance and strength groups, it was of the same direction as in the control. No statistically significant changes in SPARC levels were observed after the Bruce test – however, now the trend was decreasing. On the other hand, adiponectin showed a strong and, in strength and control, a statistically significant decrease between the timepoints 6h and 24h after the Wingate test. It remained steady throughout and after the Bruce intervention. The divergent response of SPARC and adiponectin could be attributed to the different physiological demands and stress responses induced by the Wingate and Bruce tests.

**Figure 2.**
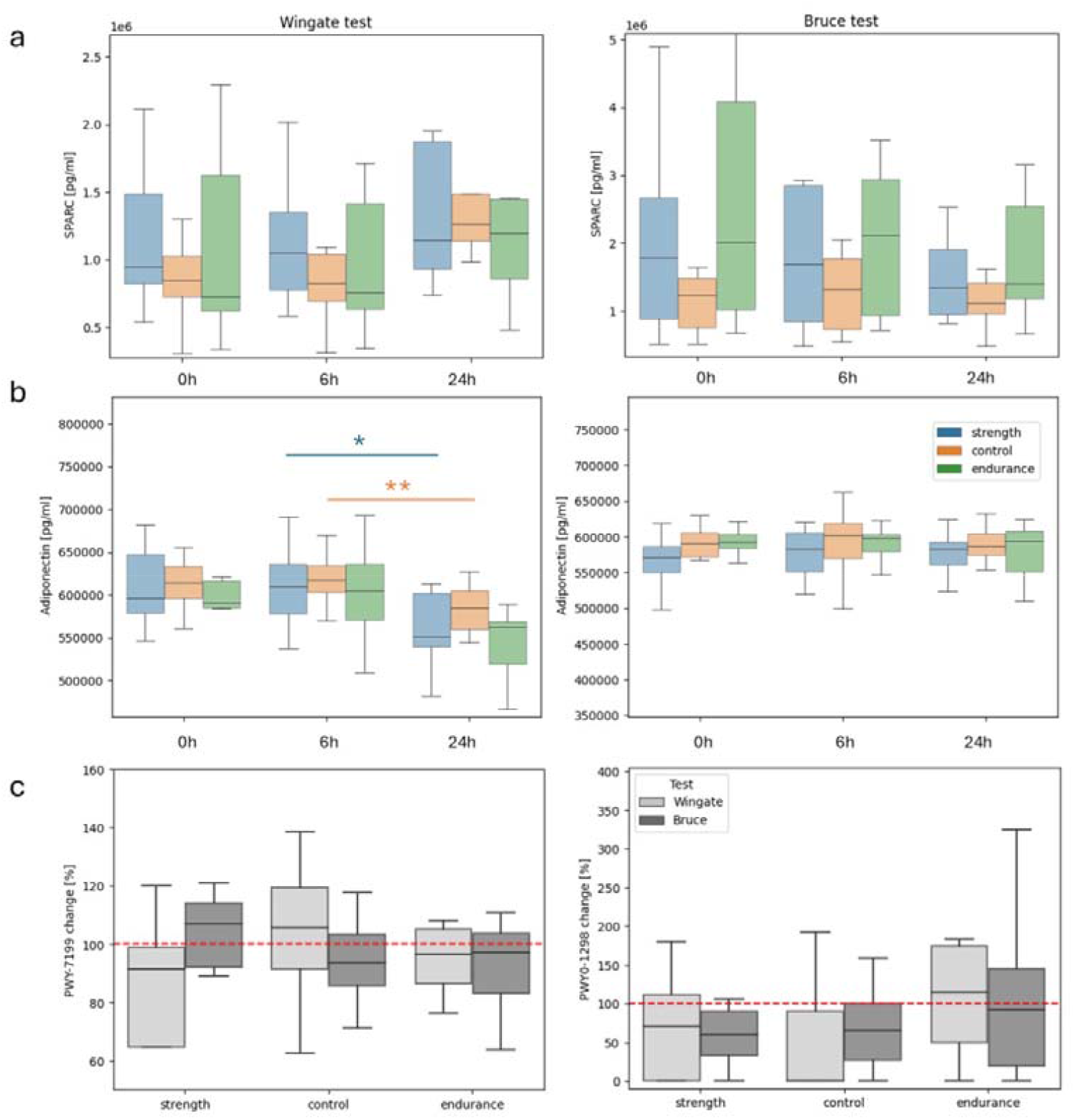
Distinct trends in the SPARC, adiponectin and pyrimidine pathways following the Wingate and Bruce tests. Different SPARC trends between timepoints before exercise, 6h after exercise and 24h after the Wingate (a) and Bruce (b) tests. Different Adiponectin trends between timepoints after the Wingate (c) and Bruce (d) tests. A relative abundance changes between timepoints 6h and 24h after the Wingate (e) and Bruce (f) interventions of the *PWY-7199: pyrimidine deoxyribonucleosides salvage pathway* and *PWY0-1298: superpathway of pyrimidine deoxyribonucleosides degradation*. A % change of 100 indicates no change; ^*^=p<0,05, ^**^=p<0,01.

We identified distinct statistically significant correlations of microbiome feature changes with the changes in adiponectin and SPARC between timepoints 6h and 24h after the Wingate and Bruce interventions. Related to the the Wingate intervention was a correlation of *PWY0-1477:_ethanolamine_utilization* with both SPARC (Superman correlation of 0.85, p<0.001, p_adjust < 0.2) and adiponectin in the strength group (r=0.90, p<0.001, p_adjust=0.05). Associated only with SPARC was *Blautia massiliensis* (r=0.83, p<0.01, p_adjust < 0.2) and the *PWY-7199: pyrimidine deoxyribonucleosides salvage pathway* (r=0.83, p<0.01, p_adjust < 0.2). Interestingly, the only association in the control group was also linked to pyrimidine deoxyribonucleosides – however, in this case it was its degradation (*PWY0-1298: superpathway of pyrimidine deoxyribonucleosides degradation* correlation with adiponectin, r=-0.92, p<0.0001, p_adjust<0.01). A further investigation revealed associations of the pathways after the Bruce test as well: *PWY-7199: pyrimidine deoxyribonucleosides salvage pathway* was correlated with TIMP1 in the endurance group (r=0.78, p<0.01, p_adjust < 0.1) and *PWY0-1298: superpathway of pyrimidine deoxyribonucleosides degradation* correlation with IL-15 in strength (r=-0.95, p<0.0001, p_adjust=0.02). Comparing relative abudance changes of the pathways between timepoints 6h and 24h after the Wingate and Bruce interventions identified nonsignificant but notable differences between the interventions and the groups (Figure 2 e,f). The results indicate a complex response of the gut microbiome to intense athletic efforts via the mechanisms of nucleotide recycling.

After the Bruce intervention, adiponectin was correlated with *PWY3O-4107: NAD salvage pathway V (PNC V cycle) and PWY-4041: γ-glutamyl cycle* (Spearman correlations of -0.92 and -0.90 respectively, both had p<0.001 and p_adjust<0.01). We identified an association of SPARC with *PWY0-1241: ADP-L-glycero-β-D-manno-heptose biosynthesis* in control (r=-0.80, p<0.01, p_adjust<0.2). On the contrary, no associations with the blood markers after the Bruce test were found in the strength group.

### Training background shapes microbiome response to exercise

#### Wingate test

The next step involved an analysis of biochemical parameters which had different trends in the training groups after the Wingate intervention. Correlations with microbiome feature abundances at timepoints 6h and 24h revealed complex associations mainly in the strength group (Table 1). This confirmed our hypothesis that exercise modality would impact individuals adapted to it the most. We found two pathways in the strength group which were positively correlated with multiple blood markers. *PWY-5989: stearate biosynthesis II* was associated with IL-15, Il-1a and Il-10 while *PWY0-1477: ethanolamine utilization* with adiponectin and SPARC (aforementioned in the previous section). In addition, we identified a number of blood marker connections with *Blautia spp*., *Lactococcus lactis, Alistipes onderdonkii, Sutterella wasdworthensis* and *Roseburia intestinalis*, as well as the *PWY-6123: inosine-5’-phosphate biosynthesis* I and *GLUDEG-I-PWY: GABA shunt pathways*.

**Table 1.** Spearman correlations of microbiome and parameter changes between timepoints 6h and 24h after the Wingate intervention.

| Group | Parameter | Microbiome feature | Spearman correlation | Adjusted p-value |
| --- | --- | --- | --- | --- |
| Control | Adiponectin [pg/ml] | PWY0-1298: superpathway of pyrimidine deoxyribonucleosides degradation | -0.92 | **<0.01** |
|  | TIMP1 [pg/ml] | Parasutterella excrementihominis | -0.87 | **0.04** |
|  | Resistin [pg/ml] | PWY-6305: superpathway of putrescine biosynthesis | 0.86 | **<0.01** |
| Strength | Adiponectin [pg/ml] | PWY0-1477: ethanolamine utilization | 0.90 | <0.1 |
|  | IL-15 [pg/ml] | PWY-5989: stearate biosynthesis II (bacteria and plants) | 0.90 | <0.1 |
|  | IL-1a [pg/ml] | PWY-5989: stearate biosynthesis II (bacteria and plants) | 0.90 | <0.1 |
|  | Resistin [pg/ml] | Blautia glucerasea | 0.88 | <0.1 |
|  | IL-10 [pg/ml] | PWY-6123: inosine-5'-phosphate biosynthesis I | 0.86 | <0.1 |
|  | IL-10 [pg/ml] | PWY-5989: stearate biosynthesis II (bacteria and plants) | 0.86 | <0.1 |
|  | FSTL1 [pg/ml] | Lactococcus lactis | 0.86 | <0.2 |
|  | SPARC [pg/ml] | PWY0-1477: ethanolamine utilization | 0.85 | <0.2 |
|  | SPARC [pg/ml] | Blautia massiliensis | 0.83 | <0.2 |
|  | Resistin [pg/ml] | GGB9758_SGB15368 | 0.82 | <0.2 |
|  | Resistin [pg/ml] | Alistipes onderdonkii | 0.82 | <0.2 |
|  | SPARC [pg/ml] | PWY-7199: pyrimidine deoxyribonucleosides salvage | 0.82 | <0.2 |
|  | Resistin [pg/ml] | Candidatus_Cibiobacter_qucibialis | 0.82 | <0.2 |
|  | Resistin [pg/ml] | Ruminococcaceae_unclassified_SGB15234 | 0.82 | <0.2 |
|  | IL-10 [pg/ml] | GLUDEG-I-PWY: GABA shunt | -0.85 | <0.2 |
|  | Resistin [pg/ml] | Sutterella wadsworthensis | -0.86 | <0.2 |
|  | LIF [pg/ml] | Roseburia intestinalis | -0.87 | <0.2 |
| Endurance | LIF [pg/ml] | Faecalibacterium_SGB15346 | 0.93 | <0.1 |
|  | TIMP1 [pg/ml] | Bacteroides uniformis | -0.92 | <0.2 |
|  | TIMP1_[pg/ml] | Bacteroides_ovatus | -0.90 | <0.2 |
|  | TIMP1_[pg/ml] | PWY0-1261:_anhydromuropeptides_recycling_l | -0.90 | <0.2 |

On the other hand, we observed an insignificant upward trend of the IL-10 marker in the control group between the timepoints 6h and 24h after the Wingate test. The levels of this marker would decrease in the other groups – significantly in endurance (Figure 3a). IL-10 is an anti-inflammatory cytokine that regulates immune response. Its decrease indicates stress, resulting from physical exertion, which in the short term may reduce the individual’s anti-inflammatory abilities. We found no microbiome correlations with Il-10 in the control group after the Wingate intervention.

**Figure 3.**
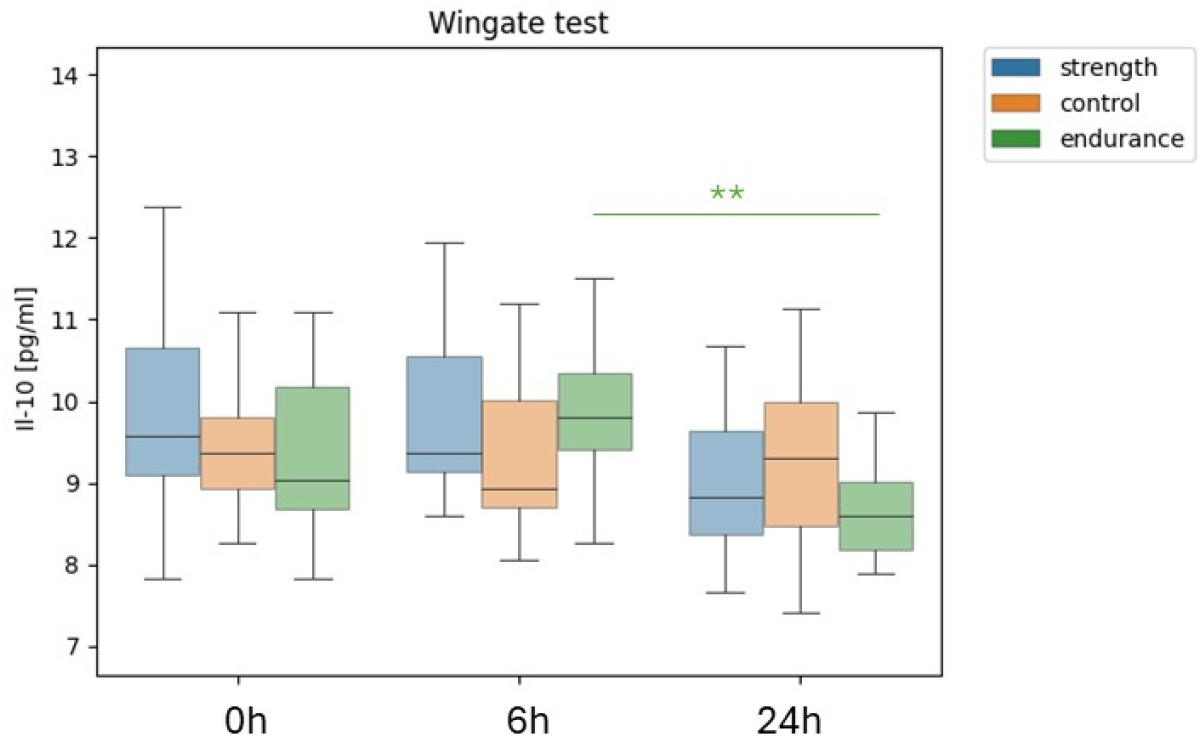
Differences in IL-10 levels between training groups after the Wingate test. Average IL-10 concentration at all timepoints in control, strength and endurance groups after the Wingate test; ^*^=p<0,05, ^**^=p<0,01.

#### Bruce test

When we analysed biochemical parameter differences after the Bruce intervention, we found complex connections mainly in the endurance group, which is adapted to this kind of efforts the most (Table 2). The majority of correlations were linked to TIMP1 and related to amino acid metabolism, carbohydrate metabolism and bacterial biosynthesis. We found 2 species and 2 pathways which were significantly associated with at least 2 blood markers: *Eubacterium rectale, Blautia wexlerae, PWY-5188:tetrapyrrole biosynthesis I from_glutamate* and *PWY-5030: L-histidine degradation III*. Notably, all of them were linked to TIMP1 and TfR, while *Eubacterium rectale* was additionally associated with Il-15.

**Table 2.**
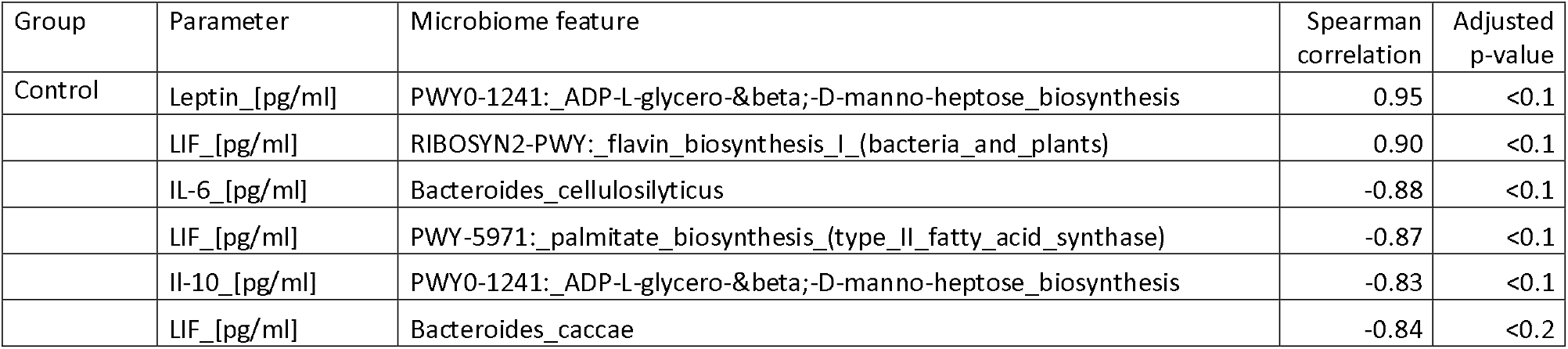
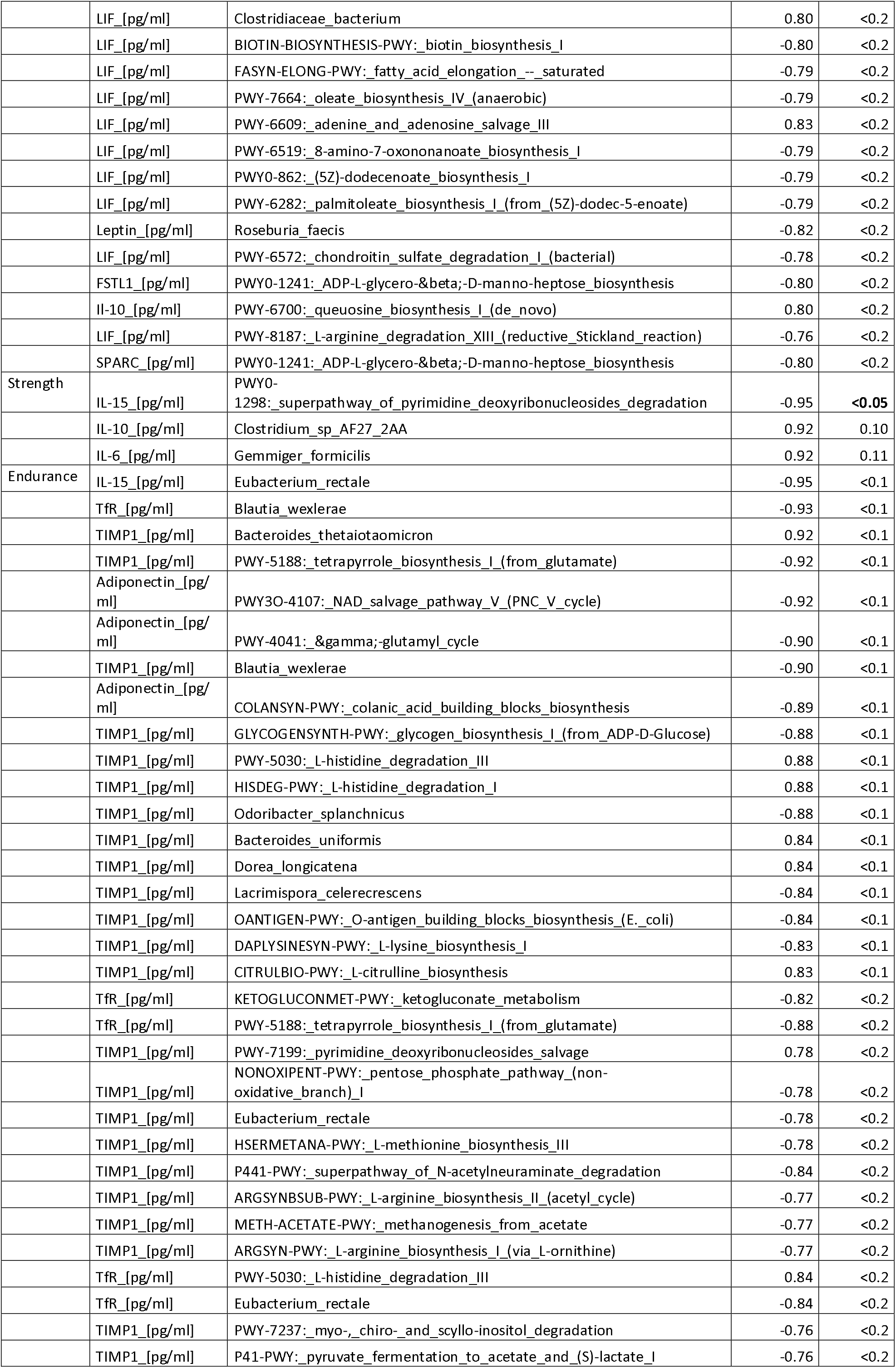
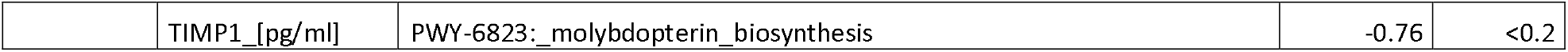
Spearman correlations of microbiome and parameter changes between timepoints 6h and 24h after the Bruce intervention.

There was also a number of associations within the control group. This time, however, most correlations were with LIF - particularly those involved in fatty acid biosynthesis, biotin biosynthesis, and amino acid metabolism. The *PWY0-1241: ADP-L-glycero-β-D-manno-heptose biosynthesis* pathway was the microbiome parameter linked to multiple different blood markers: leptin (Spearman correlation 0.95, p_adjust <0.1), FSTL1 (r=-0.80, p_adjust <0.2) and SPARC (r=-0.80, p_adjust <0.2).

Following the observation that the strength group had substantially fewer connections than the other groups, we noticed a different trend of the two parameters, namely IL-1a and TIMP1, in the strength group after the Bruce test (Figure 4). The trend of IL-1a between timepoints 6h and 24h would rapidly decrease in control and endurance, while the mean of the parameter in the strength group appeared to experience a slight increase. Similarly, TIMP1 climbed rapidly between timepoints 6h and 24h in the strength group, while it appeared steadier in the other two groups. Both IL-1a and TIMP1 are markers of response to physical exertion. Their decrease indicates an expected inflammatory behaviour and initiation of body repair mechanisms. A different response in the strength group adapted to an anaerobic type of training (further referred to as the nonresponders) and control or endurance (further referred to as the responders) might indicate greater stress and an altered response to the aerobic exercise in the strength group.

**Figure 4.**
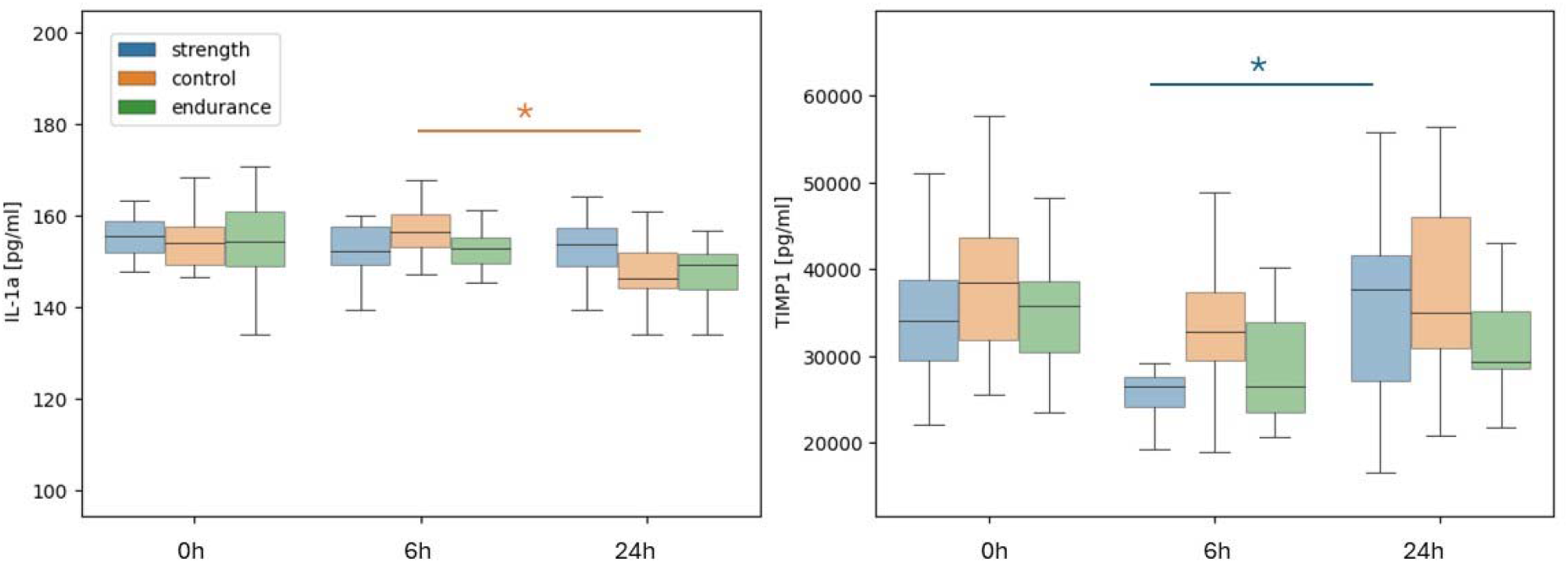
Differences in IL-1a and TIMP1 levels between training groups after the Bruce test. IL-1A (a) and TIMP1 (b) trends after the Bruce test show a different response of the strength group. ^*^=p<0,05.

### Baseline microbiome composition can determine adaptation response to the aerobic intervention

We wondered whether the inhibited response to the Bruce intervention in the strength group could be associated with the microbiome composition at baseline before the intervention. Differential enrichment analysis with Maaslin2 between the strength group (nonresponders) and control combined with the endurance group (responders) resulted in the identification of two species, *Clostridium phoceensis* and *Catenibacterium spp AM22_15*, which were enriched in nonresponders versus responders at baseline before the intervention (Figure 5). A function-based enrichment analysis identified pathways related to guanosine and adenosine biosynthesis as well as pyrimidine nucleobases salvage, however, the results did not pass false discovery correction.

**Figure 5.**
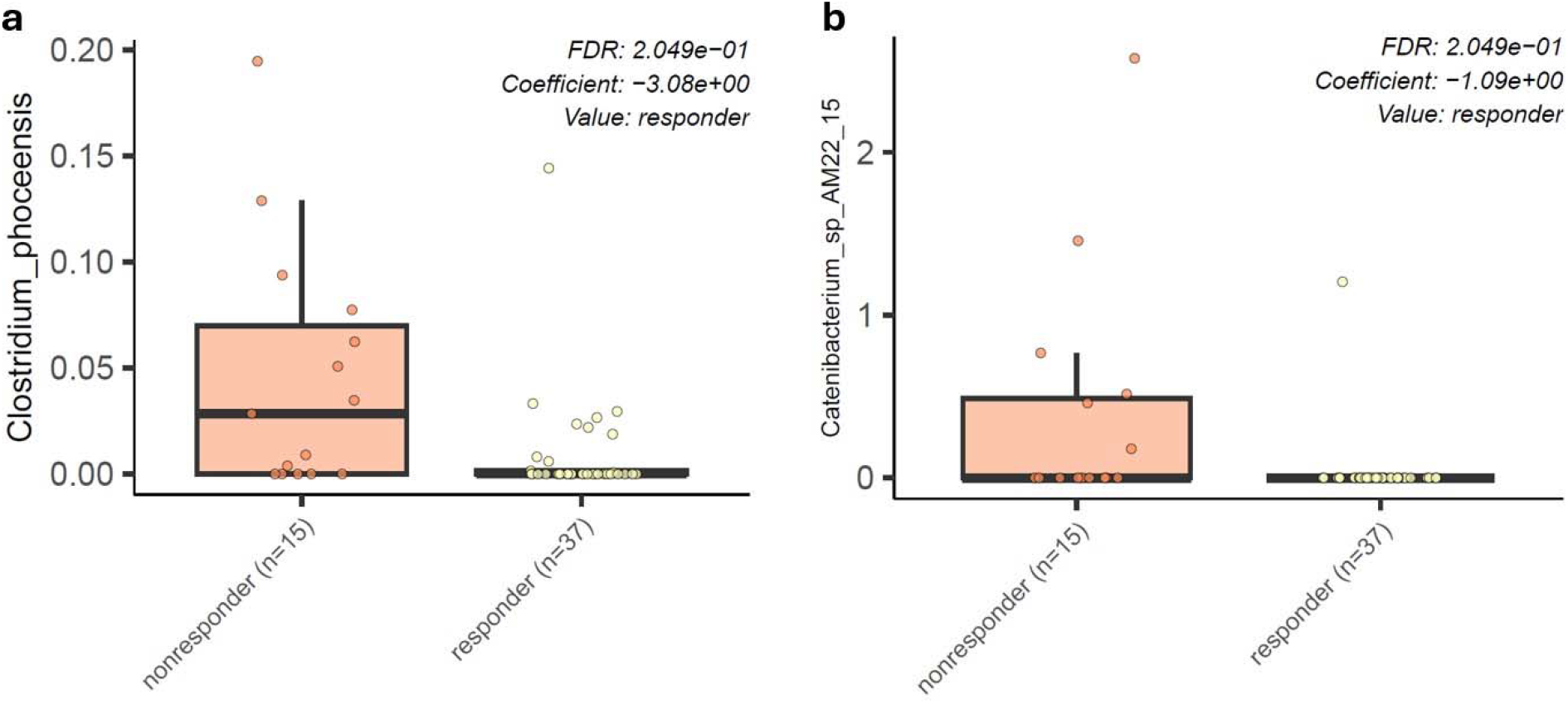
Baseline enrichment of specific bacterial species in Bruce test non-responders. Differential enrichment analysis identified two species that were more enriched in the baseline microbiome of the nonresponder group to the Bruce test.

## Discussion

Our study is one of the first to combine metagenomics and blood plasma markers in order to enhance the understanding of the role the gut microbiome plays in recovery from a training intervention. Overall, our results reveal complex physiological and biochemical responses elicited by different exercise modalities and linked to training backgrounds. The multi-omics approach, combined with a longitudinal study design, resulted in the identification of inter-group signals not discovered when investigating metagenomic data alone. In addition, it allowed us to hypothesize about potential mechanisms through which the gut microbiome modulates recovery from acute training interventions.

The changes in biochemical parameters following the Wingate and Bruce tests highlight distinct physiological responses to different exercise modalities. Changes appeared mostly between the 6^th^ and 24^th^ hour after the intervention. OSM levels significantly decreased across all groups post-exercise, indicating a general reduction in inflammation consistent with muscle regeneration processes [23]. Similarly, leptin levels showed a decrease (significant in the strength group) after the Wingate test, reflecting the hormone’s sensitivity to anaerobic exercise and its role in energy regulation [24]. Follistatin-like 1 (FSTL1) showed a biphasic response, with an initial increase followed by a significant decrease, highlighting its involvement in inflammation and muscle repair after both types of exercise [25]. Markers such as LIF, resistin, IL-6 and TfR remained stable in both tests, indicating that these pathways were not significantly affected by the exercise protocols used.

SPARC and adiponectin were the only parameters which were conserved across control, strength and endurance but experienced a different trend as a result of the two interventions. We noted connections of the markers with two pathways related to pyrimidine deoxyribonucleosides salvage and degradation. These pathways can influence adiponectin levels through several indirect mechanisms. Metabolic pathways related to nucleotide degradation affect the levels of metabolites associated with oxidative stress and inflammation, which are known factors that lower adiponectin levels [26],[27]. Nucleotides play a key role in cellular energy, and their degradation can affect energy availability, which in turn influences adiponectin function in regulating energy metabolism and insulin sensitivity [28],[29]. Additionally, nucleotide degradation products can affect cellular signalling, altering inflammatory and metabolic processes and impacting adiponectin levels [30], [31]. The observed changes in adiponectin levels post-exercise may also be influenced by the body’s prioritization of nucleotide degradation pathways during recovery. Acute high-intensity exercise, such as the Wingate test, likely increases the demand for nucleotide catabolism, which is essential for DNA repair and energy production—critical processes during the recovery phase [32],[33],[34],[35]. This prioritization of nucleotide metabolism could impact the synthesis or regulation of other proteins and hormones, including adiponectin, which is involved in lipid metabolism, insulin sensitivity, and anti-inflammatory responses [36]. In addition, nucleotide degradation products have been shown to influence cellular signalling pathways that regulate inflammation and metabolism [37], which may further modulate adiponectin levels. The decreased adiponectin levels observed after the Wingate test may reflect a shift in metabolic priorities, favouring energy production and cellular repair processes [38].

This divergent response of adiponectin could also be attributed to the different physiological demands and stress responses induced by the Wingate and Bruce tests. The Wingate test, which is a high-intensity anaerobic exercise, may induce a more pronounced acute inflammatory response, thereby affecting adiponectin levels more than the aerobic nature of the Bruce test, which typically induces a milder systemic response. This observation suggests that the type and intensity of exercise play a critical role in modulating adiponectin levels and highlights the importance of considering exercise-specific responses in studies of metabolic and inflammatory markers [27],[26],[39]. The pyrimidine deoxyribonucleoside super pathway involves the degradation of pyrimidine nucleotides, which are essential components of DNA and RNA. The activity of this pathway may be linked to cellular and metabolic stress responses, potentially influencing systemic inflammation and metabolic regulation [30],[35],[40].The strong negative correlation between this pathway and adiponectin suggests that higher pyrimidine catabolic activity may be associated with lower adiponectin levels. This may be due to increased metabolic demands and cellular turnover during high-intensity exercise, such as the Wingate test, leading to changes in systemic markers such as adiponectin. It is possible that processes such as energy production and DNA repair are prioritized over the production of adiponectin in the immediate aftermath of exercise, leading to the observed changes. Understanding these biochemical pathways and their interactions with metabolic markers may provide deeper insights into the molecular mechanisms underlying exercise-induced metabolic adaptations in different training groups [28],[41].

Following the Bruce test, an aerobic exercise protocol, we identified two bacterial species, *Eubacterium rectale* and *Blautia wexlerae*, and two metabolic pathways, *PWY-5188 (tetrapyrrole biosynthesis I from glutamate)* and *PWY-5030 (L-histidine degradation III)*, that were significantly associated with at least two blood markers. Notably, all were associated with TIMP1 (Tissue Inhibitor of Metalloproteinases 1) and TfR (Transferrin Receptor), while *Eubacterium rectale* was additionally associated with IL-15. *Eubacterium rectale* and *Blautia wexlerae* are known butyrate-producing bacteria that play a critical role in maintaining gut health and influencing systemic inflammation [42]. Butyrate has been shown to have anti-inflammatory properties and can modulate immune responses [43], which may explain the association of these bacteria with TIMP1 and TfR. TIMP1 is involved in tissue remodeling and has been linked to muscle recovery processes [44], while TfR plays a role in iron metabolism, which is critical for oxygen transport and energy production during prolonged exercise [45].

The association of *Eubacterium rectale* with IL-15, a cytokine involved in muscle maintenance and immune responses, suggests a potential link between gut microbiome and muscle physiology. IL-15 is known to be upregulated following endurance exercise and plays a role in promoting muscle growth and enhancing immune function [46]. IL-15 also seems to play a role in reducing adipose tissue mass [46]. This finding suggests the possibility of a gut-muscle axis, in which specific gut bacteria modulate exercise-induced muscle adaptations through their metabolic activities.

The *tetrapyrrole biosynthesis pathway (PWY-5188)* and *L-histidine degradation pathway (PWY-5030)* provide further insight into the metabolic changes that occur in response to endurance exercise. The tetrapyrrole biosynthesis pathway is crucial for heme production, which is necessary for oxygen transport and mitochondrial respiration—processes that are particularly important during sustained aerobic exercise [47]. The L-histidine catabolic pathway involves the breakdown of histidine, which can affect histamine production and ammonia detoxification. Both processes are relevant in the context of prolonged exercise, where histamine is involved in vasodilation and immune response, and ammonia detoxification is critical to prevent metabolic acidosis [48].

These findings suggest that the specific gut microbiome and metabolic pathways associated with endurance exercise may contribute to the modulation of physiological adaptations and recovery processes through their effects on inflammation, iron metabolism, and energy production. Further research is needed to understand the precise mechanisms by which these bacteria and pathways interact with host physiology and to explore potential interventions that could enhance recovery and performance in endurance exercise.

Finally, identified two species, specifically *Clostridium phoceensis* and *Catenibacterium spp AM22_15*, which were enriched in individuals with inhibited response (strength group) to the Bruce intervention at baseline. *Clostridium phoceensis* contains machinery to synthesize tryptophan and has previously been shown to respond to an acute moderate-intensity exercise intervention [49]. Tryptophan metabolism is known to play a role in immune regulation and inflammation. Alterations in tryptophan metabolism can affect the production of metabolites such as kynurenines, which have been implicated in the modulation of inflammation and immune responses [50]. The presence of *Clostridium phoceensis*, which has tryptophan-synthesizing capabilities, may suggest that the strength group has a microbiome composition that affects tryptophan metabolism, potentially leading to an altered inflammatory and stress response to aerobic exercise. *Catenibacterium spp AM22_15* was identified as enriched in the strength group at baseline before the Bruce intervention in our previous study [15]. The enrichment of these species may also contribute to the unique metabolic and inflammatory responses observed in the strength group. The distinct microbiome composition in the strength group, characterized by these specific bacterial species, may underlie the group’s inhibited response to the Bruce intervention, highlighting the need for further research into the interplay between microbiome composition, tryptophan metabolism, and exercise response.

The greatest limitation of this study is a small sample size. As discussed in our previous publication, despite the participants being athletic or untrained, the group overall was quite homogeneous. Further studies with greater sample sizes are required to validate the signals reported in this manuscript.

## Conclusions

This study highlights the complex interplay between exercise modality, training background, and the gut microbiome in shaping physiological responses. By integrating metagenomic data with blood serum markers, we observed that both aerobic and anaerobic exercise induce specific biochemical changes influenced by the gut microbiome.

Key findings include distinct responses of SPARC and adiponectin levels depending on exercise modality, suggesting that these markers are differentially modulated by aerobic and anaerobic efforts. In strength-trained individuals, unique associations between specific gut microbes and blood markers were observed after the anaerobic Wingate test, suggesting that training background influences microbiome-mediated responses to exercise. Conversely, endurance athletes showed correlations between butyrate-producing bacteria and markers related to muscle recovery and iron metabolism after the aerobic Bruce test, suggesting a potential gut-muscle axis.

A baseline enrichment of certain bacterial species, such as *Clostridium phoceensis* and *Catenibacterium spp*., in strength athletes was associated with an inhibited response to the aerobic intervention. This suggests that initial microbiome composition may influence how individuals adapt to different types of exercise.

These findings highlight the important role of the gut microbiome in modulating exercise-induced physiological adaptations. Understanding these relationships could lead to personalized microbiome-targeted interventions to enhance athletic performance and recovery based on individual training backgrounds.

## Data Availability Statement

The data supporting the findings of this study are openly available in the European Nucleotide Archive (ENA) at https://www.ebi.ac.uk/ena/, reference number PRJEB60692.

## Institutional Review Board Statement

Approved by the Bioethics Committee for Clinical Research of the Regional Medical Society in Gdansk (KB-27/18).

